# Structure determination from lipidic cubic phase embedded microcrystals by MicroED

**DOI:** 10.1101/724575

**Authors:** Lan Zhu, Guanhong Bu, Liang Jing, Dan Shi, Tamir Gonen, Wei Liu, Brent L. Nannenga

## Abstract

The lipidic cubic phase (LCP) technique has proved to facilitate the growth of high-quality crystals that are otherwise difficult to grow by other methods. Because crystals grown in LCP can be limited in size, improved techniques for structure determination from these small crystals are important. Microcrystal electron diffraction (MicroED) is a technique that uses a cryo-TEM to collect electron diffraction data and determine high-resolution structures from very thin micro and nanocrystals. In this work, we have used modified LCP and MicroED protocols to analyze crystals embedded in LCP. Proteinase K in LCP was used as a model system, and several LCP sample preparation strategies were tested. Among these, treatment with 2-Methyl-2,4-pentanediol (MPD) and lipase were both able to reduce the viscosity of the LCP and produce quality cryo-EM grids with well-diffracting crystals. These results set the stage for the use of MicroED to analyze other microcrystalline samples grown in LCP.

## INTRODUCTION

Structural determination of membrane proteins has been difficult primarily due to their low expression and low stability once isolated from their native membranes. Despite these difficulties, the number of membrane protein crystal structures has surged in recent years due to multiple technical breakthroughs, including the lipidic cubic phase (LCP) technique, which provides a lipid environment close to that of the native membrane protein environment (*1*). Since the high-resolution structure from LCP was determined using bacteriorhodopsin in 1997 (*2, 3*), there are now over 120 unique membrane proteins structures at atomic resolution have been obtained in LCP (*4–6*). However, one challenge with this technique is that when membrane proteins crystallize in the LCP, it is often in the form of very small microcrystals that are too small to withstand the radiation damage used during synchrotron X-ray diffraction data collection. Serial femtosecond crystallography (SFX), which utilizes ultra-fast x-ray pulses to capture diffraction from these LCP embedded membrane protein microcrystals before they are destroyed by radiation damage, has recently been employed with great success (*7*). Nevertheless, the LCP-SFX technique is very time and resource intensive and is not widely accessible with only a few operational worldwide X-ray free electron laser facilities thus far(*8*). Moreover, this technology requires a very large number of microcrystals to collect enough diffraction patterns required to construct a complete dataset for structure determination. Therefore, to improve the study of important integral membrane proteins and uncover new high-resolution details in a more high-throughput fashion, new methods for structure determination need to be developed for the small crystals grown in LCP.

The advent of a new technique, microcrystal electron diffraction (MicroED) in 2013, offers an alternative for the structure determination of proteins from microcrystal samples (*9*). MicroED is a method that is used to collect electron diffraction patterns from sub-micron sized three-dimensional crystals in the electron microscope (*10, 11*). Microcrystals are deposited on electron microscopy grids followed by sample blotting, as electrons cannot penetrate thick samples. To ensure the sample remains hydrated in the vacuum of the electron microscope and for reduced radiation damage, the samples are vitrified and kept at cryogenic temperatures. The continuous rotation data collection strategy for MicroED allows multiple diffraction patterns taken from one single crystal with extremely low electron dose, resulting in a series of diffraction patterns that can be indexed, integrated, and processed with crystallographic data processing software without any prior knowledge of unit cell parameters or geometry(*12*). Since the initial implementation of MicroED, there have been further efforts in improving this method to determine biomolecular structures (*10*).

In this work, our goal is to combine MicroED with LCP microcrystallography methods (LCP-MicroED) to determine structures from microcrystals within the LCP matrix. Here, we report the first MicroED structures of a model protein, Proteinase K, that has been embedded in LCP. This sample was chosen because it has served as a model sample for both MicroED and new LCP-based serial crystallography methods(*13–16*). By treating the LCP samples with different reagents to lower the viscosity of the LCP samples, we identified two strategies – dilution using MPD and treatment of the sample with lipase – that led to high-quality MicroED samples. Both strategies were used on the LCP-proteinase K samples to successfully determine the structure of proteinase K to 2.0 Å resolution using MicroED.

## RESULTS

Due to the intrinsic high viscosity, LCP is not well-suited to standard MicroED sample preparation protocols (*17*). In order to identify new sample preparation conditions and protocols that would allow MicroED to be used on samples embedded in LCP, we chose a well-studied model system, Proteinase K, which has previously been used benchmark both new LCP-based diffraction (*18*) and MicroED methods (*14, 19*). Proteinase K microcrystals were grown in batch and reconstituted into the LCP matrix to be used for further studies on LCP sample preparation for MicroED.

### LCP sample conversion for EM grid deposition

We initially focused on the identification and optimization of sample preparation conditions that would allow the collection of MicroED data from crystals embedded or grown in LCP. Because the viscosity of LCP matrix is too high to be directly deposited on EM grid and effectively blotted thin enough to be penetrated by the electron beam, we introduced two different strategies to reduce the viscosity: 1) by mixing with cryoprotectants (also known as spongifiers) to convert into a liquid analogue of cubic phase, also called sponge phase (*20*); 2) treatment with lipase to hydrolyze matrix lipids and convert LCP into a two-liquid phase system of water/glycerol solution and oleic acid (*21*).

First, we screened the following cryoprotectants at various concentrations: 2-Methyl-2,4-pentanediol (MPD), PEG200, and PEG400 (*22, 23*). These three agents are commonly used in traditional protein crystallization screening and cryoprotection solutions. Initial tests were conducted with blank LCP by mixing of the host lipid monoolein (MO) and proteinase K precipitant buffer without protein. All three cryoprotectant additives could convert the LCP to a less viscous sponge phase in syringe mixing, in the range of 6-18% with MPD, 24-40% with PEG200, and 32-48% with PEG400, respectively, which are consistent with previous sponge phase transition studies (*23*).

We then tested the absorption of the formed sponge phase on blotting paper, in order to determine how well these converted LCP samples could be blotted in order to generate a thin sample for MicroED. In this step, samples were expelled out of the mixing syringe and deposited on the blotting paper, without external blotting force applied. Compared with the LCP droplet (Figure 1a), which retains its shape and does not blot on the filter paper, the three cryoprotectant-converted sponge phase samples showed absorption on the filter paper (Figure 1b-d). MPD, which has the lowest viscosity of the solutions tested (approximately 34 cP compared with 48 cP and 92 cP for PEG200 and PEG400, respectively (*24*)), as well as the lowest final ratio added to the LCP (6-18%), shows the most significant blotting relative to the PEGs. When similar blotting tests were conducted on EM grids, the MPD-converted sponge phase samples produced the most consistently thin samples for successful TEM visualization. Therefore, MPD was selected for further experiments with proteinase K crystals embedded in LCP.

**Figure 1.**
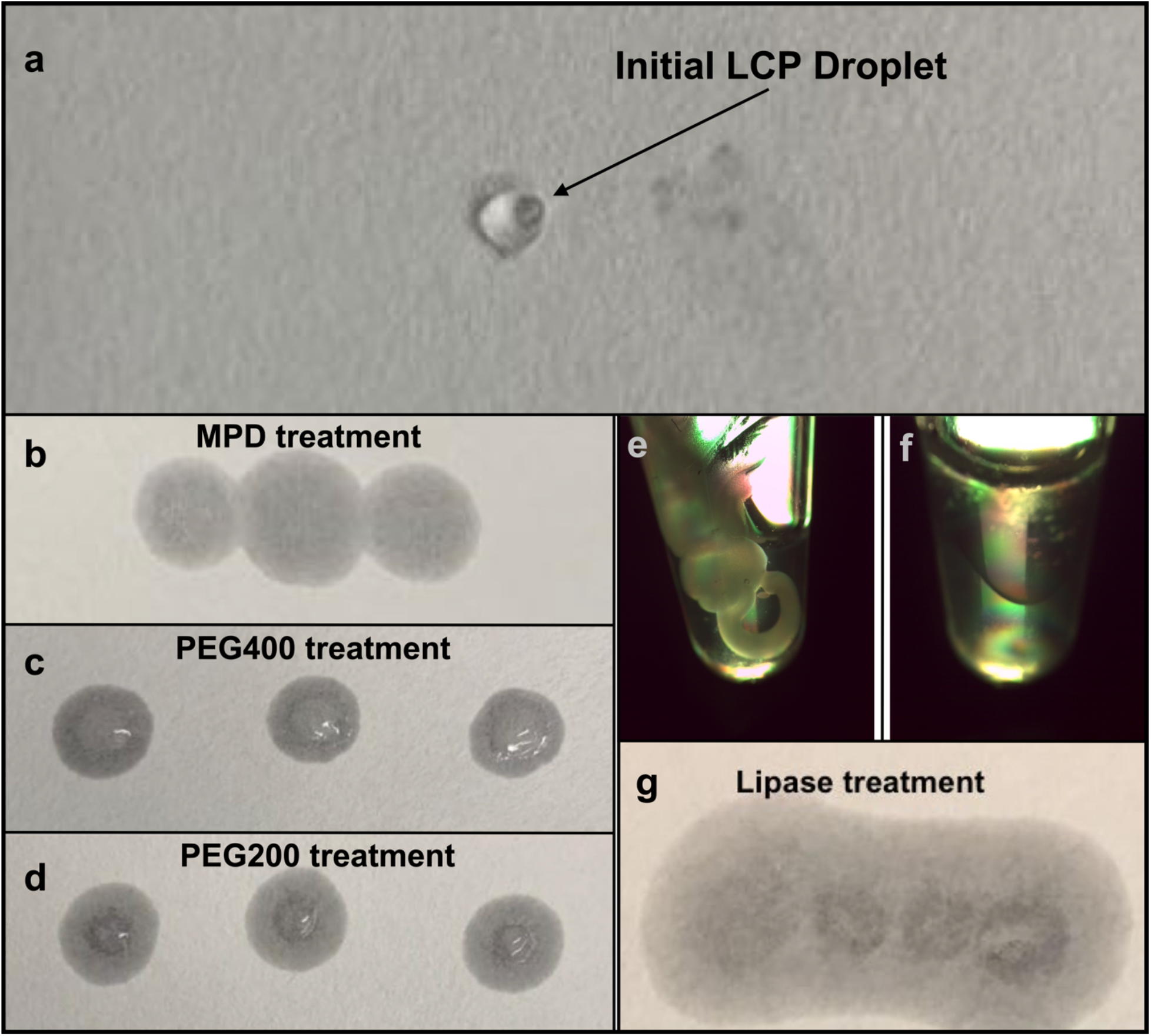
LCP phase conversion by the addition of spongifiers or lipase treatment to generate low-viscosity liquid-like sample suitable for MicroED grid preparation. a) highly viscous LCP sample is not able to be absorbed into the blotting paper, but rather retains its shape. b-c) the treatment with the phase converting buffers supplemented with three spongifiers, MPD (b), PEG400 (c), and PEG200 (d), respectively, converted the LCP sample to a liquid-like phase, which penetrated the blotting paper. e-g) the lipase hydrolysis treatment of LCP sample to form two liquid phases, e) the LCP stream (white solid stream in the tube) with freshly prepared lipase solution mixed in a ratio of 1:1 (v/v) before treatment, f) the LCP sample was separated into two liquid phases after 14-hour treatment; g) the lipase hydrolyzed sample penetrated the blotting paper indicating it can be blotted from the surface of an EM grid.

In addition to the treatment with sponge phase inducing additives, we investigated an alternative enzyme hydrolysis method by treating LCP with lipase to hydrolyze host lipid molecules and transition the cubic phase to a two-liquid phase system (*21*). Blank LCP was again used to find an optimal hydrolysis ratio of LCP with lipase, as well as the minimum treatment time to completely separate LCP into two liquid phases. LCP sample was expelled from the syringe mixer into a 0.2 mL microfuge tube, and freshly prepared lipase solution at 50 mg/mL was directly added on top of LCP sample without additional pipetting. It was found that after a 14-hour treatment in a 1:1 ratio of LCP and freshly prepared lipase solution incubated at 20 °C, the solid cubic phase (Figure 1e) was completely separated into two liquid layers (Figure 1f). This lipase hydrolyzed sample also penetrated the blotting paper without any visible LCP residue on the surface (Figure 1g). As with the MPD treated samples described above, this strategy was then tested with LCP-proteinase K crystal samples to study crystal survival and grid preparation for data collection.

### Microcrystal survival during LCP conversion

We then followed the batch crystallization method to grow proteinase K microcrystals in solution (Figure 2a) and reconstituted them into LCP by dual-syringe mixing with the host lipid monoolein in a lipid:crystal solution ratio of 3:2 (v/v). Proteinase K microcrystals survived after reconstituted into LCP (Figure 2b) and were used for phase conversion test with MPD or lipase treatment.

**Figure 2.**
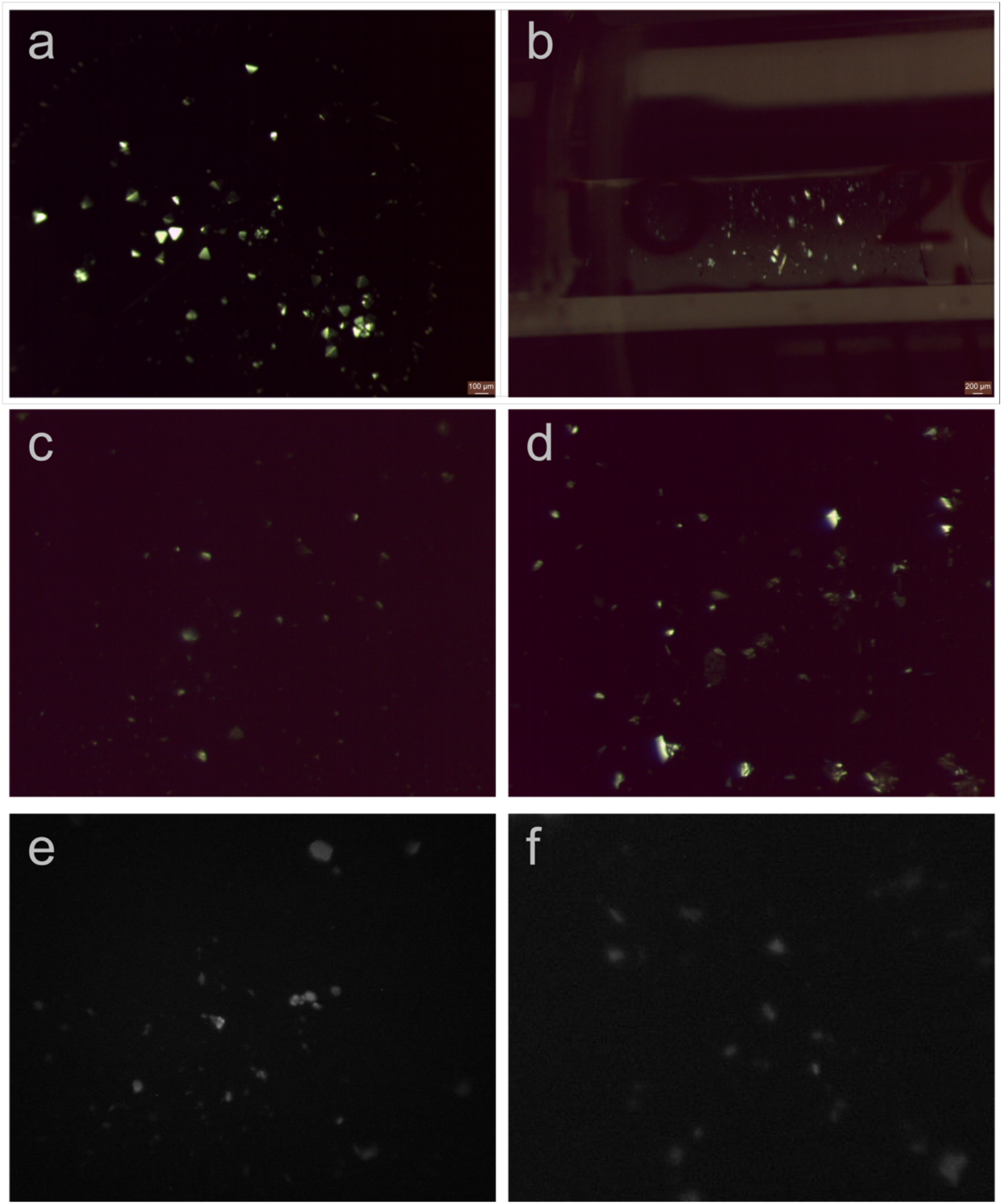
Proteinase K microcrystals were imaged before and after LCP phase conversion. Microcrystals of proteinase K grown in batch method (a) and reconstituted into the LCP matrix by syringe mixing (b), viewed with cross polarized light. Proteinase K microcrystals embedded in LCP survived after the LCP phase conversion by the converting buffer supplemented with 12.5% MPD (c) or LCP hydrolyzed by the lipase treatment (d), viewed with cross polarized light. LCP-proteinase K samples converted by MPD (e) and lipase treatment (f) were successfully blotted on the glow-discharged EM grids used for MicroED data collection, and when grids were viewed by UV, the presence of crystals could still be seen on the blotted grids.

For sponge phase conversion, we tested the range of 6-18% MPD supplemented to the proteinase K precipitant solution in increments of 0.5% as a phase converting buffer. This conversion buffer was then mixed with the LCP-proteinase K in the syringe. It was observed that the converting buffer supplemented with more than 12.5% MPD resulted in fewer/no proteinase K microcrystals or crystals with dissolved edges (crystal image not shown). Therefore, 12.5% MPD was chosen as the maximum concentration capable of to reducing the viscosity of the LCP sample (Figure 1b) without dissolving the embedded proteinase K crystals (Figure 2c).

Treatment of LCP with lipase (1:1 (v/v) ratio) was applied to LCP-proteinase K samples, whereas with these samples containing proteinase K crystals, additional proteinase K precipitant solution was added as a supplement to ensure the stability of the crystals after release from the LCP. Hydrolysis time was also re-monitored every 1 hour for the first 14 hours, and every 20 minutes afterward. After 18-hours treatment, the 1:1:2 (v/v/v) ratio of lipase:LCP-proteinase K:precipitant sample completely was digested and separated the LCP into two liquid layers, and the proteinase K crystals were released from the LCP matrix into the glycerol rich phase. Similar size and density of proteinase K microcrystals in the hydrolyzed liquid phase were observed compared to the LCP-proteinase K sample before treatment (Figure 2d).

With these sample preparation methods, the low-viscosity LCP-microcrystal solution is applied to the glow-discharged carbon coated EM grid, blotted with filter paper and vitrified in a method that is similar to preparing grids for non-LCP MicroED samples (*17*). MPD-induced sponge phase sample was further diluted by adding the converting buffer in a 1:1 (v/v) ratio, which generated a thin layer on EM grid where crystals could still be identified by UV microscopy (Figure 2e). Lipase treatment also produced grids with a similar level of microcrystals visible by UV (Figure 2f).

### MicroED analysis of treated LCP samples

Samples prepared by the methods described above were analyzed in the cryo-TEM to verify the sample thickness and the presence of proteinase K microcrystals. In both cases, the sample was thin enough such that regions of the grid with proteinase K microcrystals could be identified (Figure 3a and d). While both treatment strategies produced suitable samples, lipase-treated samples generally gave a thinner layer on the EM grid relative to the MPD-induced sponge phase samples.

**Figure 3.**
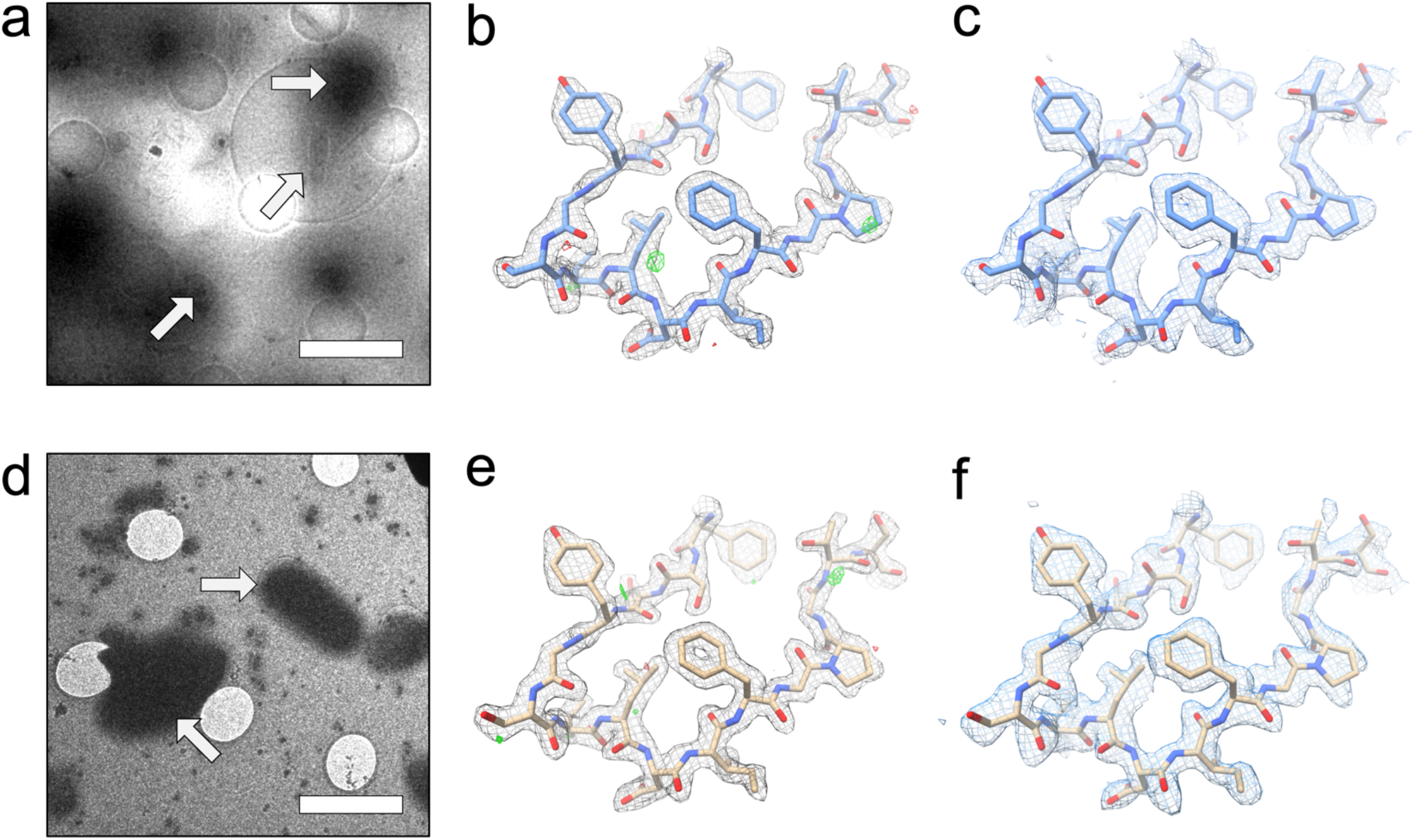
LCP-MicroED structure of proteinase K. Both MPD treated samples (a-c) and lipase treated samples (d-f) produced grids where crystals could be identified in the cryo-TEM (a - MPD and d - lipase). MicroED data collection on the crystals from both treatments produced structures at 2.0 Å. The 2F_o_-F_c_ density maps (b – MPD and e – lipase) and composite omit maps (c – MPD and f – lipase) show clear density surrounding the models. Scale bars in a and d represent 4 μm. The density maps in b and e and composite omit maps in c and f are contoured at 1.5σ and 1.0σ, respectively. The 2F_o_-F_c_ map in b and e is contoured at 3.0σ (green) and −3.0σ(red).

Standard MicroED diffraction screening, data collection, and data processing protocols (*17, 25*) were used to collect high-resolution data sets from each type of sample preparation. For the MPD converted samples, data from a total of 4 crystals were merged together to produce a final data set and a fully refined structure at 2.0 Å. In the case of the lipase digested LCP samples, data from 2 crystals were used to determine the final structure of Proteinase K, which was also at 2.0 Å resolution. Both of these methods of LCP sample preparation ultimately produced quality MicroED data and density maps and models (Table 1; Fig 3). When compared to other Proteinase K structures determined by MicroED (*15, 16, 19*), both of the LCP-proteinase K structures in this work showed similar levels of data quality (e.g. resolution, R-factors), indicating that the processing methods used did not greatly affect the crystal quality of the microcrystals. Additionally, when the final structures are compared to another MicroED proteinase K structure (PDB ID: 5I9S (*19*)), the resulting models are very similar, with all atom RMSD values of 0.59Å between the previously published structure and both the MPD and lipase treated samples. When the structures resulting from the MPD and lipase treatments are compared, the all atom RMSD between these two new structures is 0.47Å.

**Table 1.**
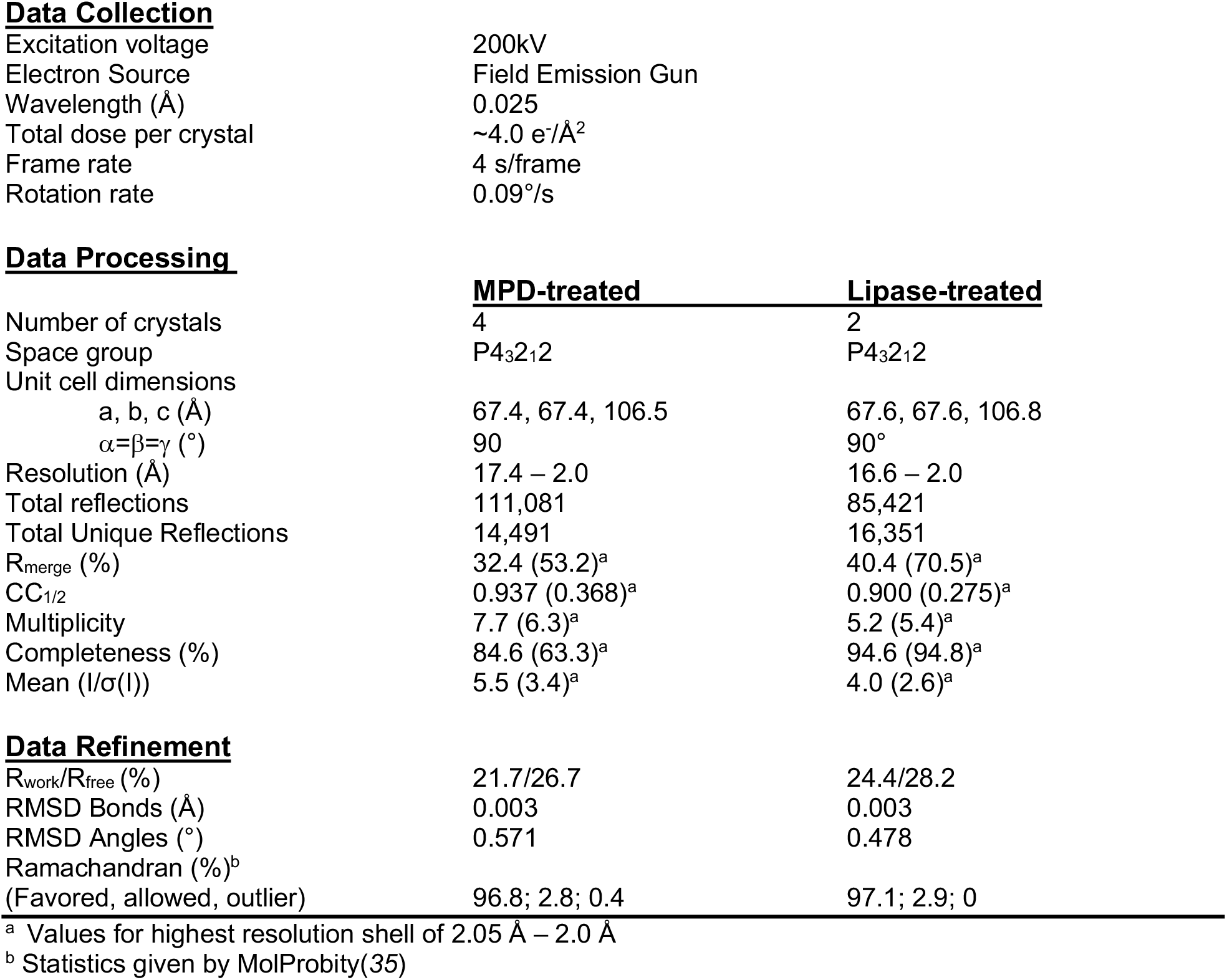
Data collection and refinement statistics

|  |  |
| --- | --- |
| Excitation voltage | 200kV |
| Electron Source | Field Emission Gun |
| Wavelength (Å) | 0.025 |
| Total dose per crystal | ~4.0 e<sup>-</sup>/Å<sup>2</sup> |
| Frame rate | 4 s/frame |
| Rotation rate | 0.09°/s |

**Data Processing**
|  | **MPD-treated** | **Lipase-treated** |
| --- | --- | --- |
| Number of crystals | 4 | 2 |
| Space group | P4<sub>3</sub>2<sub>1</sub>2 | P4<sub>3</sub>2<sub>1</sub>2 |
| Unit cell dimensions |  |  |
| a, b, c (Å) | 67.4, 67.4, 106.5 | 67.6, 67.6, 106.8 |
| α=β=γ (°) | 90 | 90° |
| Resolution (Å) | 17.4 – 2.0 | 16.6 – 2.0 |
| Total reflections | 111,081 | 85,421 |
| Total Unique Reflections | 14,491 | 16,351 |
| R<sub>merge</sub> (%) | 32.4 (53.2)<sup>a</sup> | 40.4 (70.5)<sup>a</sup> |
| CC<sub>1/2</sub> | 0.937 (0.368)<sup>a</sup> | 0.900 (0.275)<sup>a</sup> |
| Multiplicity | 7.7 (6.3)<sup>a</sup> | 5.2 (5.4)<sup>a</sup> |
| Completeness (%) | 84.6 (63.3)<sup>a</sup> | 94.6 (94.8)<sup>a</sup> |
| Mean (I/σ(I)) | 5.5 (3.4)<sup>a</sup> | 4.0 (2.6)<sup>a</sup> |

**Data Refinement**
|  |  |  |
| --- | --- | --- |
| R<sub>work</sub>/R<sub>free</sub> (%) | 21.7/26.7 | 24.4/28.2 |
| RMSD Bonds (Å) | 0.003 | 0.003 |
| RMSD Angles (°) | 0.571 | 0.478 |
| Ramachandran (%)<sup>b</sup> |  |  |
| (Favored, allowed, outlier) | 96.8; 2.8; 0.4 | 97.1; 2.9; 0 |

<sup>a</sup> Values for highest resolution shell of 2.05 Å – 2.0 Å<sup>b</sup> Statistics given by MolProbity(35)

## DISCUSSION

These results represent the first-time microcrystals embedded in LCP have yielded high-resolution structures by MicroED. This proof-of-concept study and methodology paves the way for future MicroED applications to LCP membrane protein crystallography for challenging membrane protein targets that only form micro- or nanocrystals. To expand the application of LCP-MicroED to membrane proteins that grow as very small microcrystals in LCP, further developments and optimizations based on these initial methods can be explored. Because every LCP-microcrystal sample is unique, the expansion of these reported methods into a suite of techniques will be important for its broad applicability. For the method of using spongifiers for phase conversion, a broader spectrum of spongifiers should be investigated for LCP-MicroED. Certain polar solvents or other additives, like butanediol, Jeffamine, propylene glycol (PG), and pentaerythritol propoxylate (PPO), are commonly used in membrane protein crystallization precipitant solutions, and can also be used as spongifiers. The identification of a suite of spongifiers compatible with LCP-MicroED would allow users to choose compounds already present in the crystallization precipitant solution (or ones with similar chemical properties to the components). When screening new spongifiers two critical factors, the viscosity of the compound and the overall percentage needed to convert the phase, should be kept in mind. It was found in this study that MPD behaved much better than PEGs in microcrystal blotting on EM grid, which was attributed to its lower viscosity and lower concentration (6-18%) required for sponge phase conversion. The viscosity is not a parameter which can be tuned easily; however, spongifiers typically have a wide range of concentrations capable of sponge phase conversion (*23*). While screening spongifier concentration, it is important to note that some may affect microcrystal quality at the concentrations required to trigger sponge phase formation. Therefore, for novel spongifiers, their effects on phase conversion and crystal quality should be carefully examined.

Because lipase is able to hydrolyze the lipids that make up the LCP matrix, lipase treatment generally provided thinner samples relative to MPD treatment, thereby increasing the area which was visible in the electron microscope. In membrane protein crystallization, it is common that lipid molecules may interact with membrane proteins, and some cases even play crucial roles in protein crystal packing (*26, 27*). Though it has been previously shown that the lipase treatment of bacteriorhodopsin crystals could be performed without degrading the crystals (*28*), the lipase hydrolysis treatment method may introduce some additional deleterious effects to membrane protein crystal quality. Therefore, lipase treatment should be evaluated extensively for different membrane protein targets prior to LCP-MicroED studies.

The combination of the extraordinary properties of LCP and MicroED promises to facilitate the determination of high-resolution structures of challenging targets using just a few sub-micron thick crystals. These structural studies with MicroED could open the door to the identification of new structural information by improving resolution of poorly ordered samples (*16*), determining structures with minimal radiation damage (*15*), and facilitating the modeling of charge within structures (*29, 30*). LCP-MicroED has the potential to be a robust method to solve high-resolution structures from microcrystals grown in LCP. The successful development and use of LCP-MicroED will add another important tool to the field of structural biology.

## MATERIALS AND METHODS

### Microcrystal sample preparation

Proteinase K (catalog no. P2308, Sigma) was crystallized using the batch crystallization method by mixing equal volumes of proteinase K solution at 40 mg/mL in 0.02 M MES pH 6.5 and a precipitant solution composed of 0.1 M MES pH 6.5, 0.5 M sodium nitrate, 0.1 M calcium chloride. Proteinase K microcrystals appeared after 20 min incubation at 20 °C (*13*). Microcrystals were pelleted by centrifugation at 500 g for 5 min, resuspended in the crystallization buffer, and then reconstituted into LCP by mixing with molten monoolein host lipid in a lipid:solution ratio of 3:2 (v:v) using a dual-syringe mixer until a homogeneous and transparent LCP was formed (*22*).

### LCP microcrystal sample conversion set-up

Proteinase K microcrystals embedded in LCP were then either converted by mixing with additives to achieve a less viscous lipidic sponge phase or subjected to lipase treatment to separate into two immiscible liquid phases.

LCP-proteinase K crystal sample conversion was tested with three cryoprotectant additives, MPD, PEG200, and PEG400. Sample conversion was performed by syringe mixing 15-20 times of an LCP embedded proteinase K sample in one syringe and the conversion buffer in the other syringe. The conversion buffer was made from the initial crystallization buffer supplemented with each of different additives. Each conversion buffer was optimized with a gradient additive concentration series of 10% increments. Finer additive concentration optimization followed when an initial point was identified that was capable of converting the LCP phase to a sponge phase. Once the concentration range of supplemented additive was identified, EM grid blotting experiments were conducted to investigate the capability of the additives for producing quality MicroED samples. Among those three cryoprotectant additives, only MPD-induced sponge phase sample exhibited reproducibly good quality EM grids. MPD was then focused for further optimization to ensure crystal survival during phase conversion. Once the LCP phase was converted, LCP-microcrystal samples were transferred into a microcentrifuge tube. Crystals were centrifuged and harvested from the bottom by pipetting. These samples were applied to a glass slide to monitor crystal survival by light microscopy with cross-polarized and UV light, or deposited on EM grid for analysis in the cryo-TEM.

In the case of lipase treatment, LCP-proteinase K microcrystal samples were expelled from the LCP mixing syringe into a 0.2 mL microcentrifuge tube, followed by the addition of fresh prepared lipase solution at a volume ratio of 1:1:2 (lipase solution:LCP:crystallization buffer) directly into the same tube without mixing. The lipase used is from *Candida rugosa* (catalog no. L1754, Sigma) and is prepared at a concentration of 50 mg/mL in saturated K phosphate buffer. Incubation for at least 18 hours converts the lipidic cubic phase into a two-phase system consisting of two immiscible liquids: water/glycerol and oleic acid. Proteinase K crystals partitioned into the glycerol/water phase.

### MicroED sample preparation and data collection

After MPD or LCP treatment, proteinase K microcrystals were collected by centrifugation at 500 g for 5 min. Crystal solution was then harvested by pipetting from the microfuge tube bottom into a fresh microfuge tube. For MPD samples, the crystal solution was further diluted by adding a 1:1 ratio of fresh precipitant solution supplemented with 12.5% MPD. Cryo-TEM samples were prepared by standard MicroED sample preparation procedures (*17*). Briefly, 2 μL of crystal solution was deposited on each side of a glow-discharged holey carbon EM grid (Quantifoil 2/4), and the grid was processed with a Vitrobot Mark IV (Thermo Fisher) by blotting for 12-16 s followed by vitrification by plunging into liquid ethane. Sample preparation was optimized and screened in high-throughput fashion using a Titan Krios with CETA CMOS detector (Thermo Fisher). MicroED data collection was performed by standard methods (*17*) using an FEI TF20 equipped with a F416 CMOS detector (TVIPS).

### MicroED data processing and structure determination

MicroED data collected from MPD and lipase treated LCP-proteinase K microcrystals were and integrated in iMOSFLM (*31*). For each treatment, data from multiple crystals (4 for MPD-treated and 2 for lipase-treated) were merged and scaled in AIMLESS (*32*) to create merged data sets with high completeness. Phaser (*33*) was used to perform molecular replacement using a Proteinase K search model (PDB ID: 2ID8), and the solution was refined in phenix.refine (*34*) using electron scattering factors.

### Data availability

Coordinates and structure factors were deposited in the Protein Data Bank (PDB) under the accession code 6PQ0 and 6PQ4.

## Acknowledgements

We would like to acknowledge the use of the Titan Krios at the Erying Materials Center at Arizona State University and the funding of this instrument by NSF MRI 1531991. This work was supported by the Centre for Applied Structural Discovery (CASD) at the Biodesign Institute at Arizona State University (L.Z., G.B., L.J., W.L. and B.L.N.), a Mayo Clinic ASU Collaborative Seed Grant Award (W.L.), the Flinn Foundation Seed Grant (W.L.), the STC Program of the National Science Foundation through BioXFEL (No. 1231306; L.Z., L.J. and W.L.), the National Institutes of Health grants R21DA042298 (W.L.) and R01GM124152 (W.L. and B.L.N.). The Gonen lab is supported by funds from the Howard Hughes Medical Institute.

## Author contributions

L.Z., W.L., and B.L.N designed the experiments and protocols. L.Z., G.B. and L.J. grew crystals and prepared samples for data collection. G.B., D.S., T.G. and B.L.N. participated in data collection and data analysis. L.Z. and B.L.N prepared the figures and tables for the manuscript. L.Z., W.L. and B.L.N. wrote the manuscript with contributions from all authors. All authors read and approved the final manuscript.

